# Dynamic Causal Modelling for Functional Near-Infrared Spectroscopy Using Spatial Priors Derived from Diffuse Optical Tomography

**DOI:** 10.1101/2025.10.31.685970

**Authors:** Truc Chu, Kiyomitstu Niioka, Ippeita Dan, Sungho Tak

## Abstract

Functional near-infrared spectroscopy (fNIRS) is an optical neuroimaging technique that measures brain activity by detecting changes in oxygenated and deoxygenated hemoglobin concentrations. Although methods exist for estimating causal interactions among brain regions using fNIRS data, current approaches typically rely on node locations defined at the sensor measurement level near the cortical surface. To address this limitation, this study extends dynamic causal modeling (DCM) for fNIRS by incorporating source-level locations derived from diffuse optical tomography (DOT). The proposed method was applied to an experimental dataset comprising 104 participants recorded during a Go/No-Go response inhibition task. Bayesian model selection confirmed that DCM models using DOT-informed neuronal source locations outperformed models using sensor-level locations. Furthermore, posterior means of effective connectivity parameters validated the inhibitory influence of the right inferior frontal gyrus on regions within the motor network during response inhibition. By providing precise neuronal source localization informed by statistical parametric mapping of DOT-reconstructed data, the proposed DCM approach enables more accurate inference of directed connectivity at the neuronal level from optical density measurements. Given the compact and portable nature of fNIRS systems, this method can readily be applied in realistic, naturalistic environments to investigate network-level modulation of brain responses.

## 1. Introduction

Functional near-infrared spectroscopy (fNIRS) is an optical neuroimaging technique used to monitor brain function by measuring concentration changes in oxygenated (HbO) and deoxygenated (HbR) hemoglobin (Jöbsis, 1977; Villringer et al., 1993; Ferrari and Quaresima, 2012). The basic unit of an fNIRS system consists of a source and detector pair, in which the source emits near-infrared light, typically within the wavelength range of 650 to 950 nm, from the scalp surface. Within 650–950 nm, biological chromophores absorb weakly, allowing near-infrared photons to traverse several centimeters of tissue before detection. Changes in regional hemoglobin concentration (HbO, HbR) associated with brain activity are reflected as variation in the detected light intensity across each source-detector pair (Delpy et al., 1988).

fNIRS provides several advantages for neuroscience research compared with conventional modalities such as functional magnetic resonance imaging (fMRI), positron emission tomography (PET), and magnetoencephalography (MEG). Unlike the blood oxygenation level-dependent (BOLD) signal measured by fMRI, which indirectly reflects oxygen consumption through the physiological coupling of cerebral blood flow, oxidative metabolism, and cerebral blood volume, fNIRS provides direct measurements of changes in HbO, HbR, and total hemoglobin at subsecond temporal resolution (Boas et al., 2014). Furthermore, the compact, portable, and quiet nature of fNIRS makes it especially suitable for clinical applications, particularly in situations where other modalities are impractical, such as with individuals who have implanted electronic devices (Wheelock et al., 2019). fNIRS is also well-suited for studies involving infants (Ferradal et al., 1991; White et al., 2012), patients with stroke or seizures (Singh et al., 2014; Chalia et al., 2016), and investigations of auditory and language systems (Eggebrecht et al., 2014).

Brain activity measured with fNIRS can be characterized not only by regional activation at individual sensors but also the interactions among regions, described as either functional or effective connectivity (Yücel et al., 2021, Tak et al., 2014) Functional connectivity reflects temporal correlations in the hemodynamic signals, whereas effective connectivity reveals directed, causal influences between brain regions during cognitive processing (Friston 1994, 2011). Dynamic causal modelling (DCM) is a method for estimating effective connectivity within a Bayesian framework (Friston et al., 2003) and has been adapted for fNIRS data (Tak et al., 2015). In this method, a generative model of the optical signal is coupled to hemodynamic and neuronal state equations, and Bayesian inversion yields posterior parameter distributions. The DCM for fNIRS method has been validated in a healthy infant, showing that the effective connectivity estimates from fNIRS closely match those obtained from simultaneously recorded fMRI (Bulgarelli et al., 2018).

However, the existing DCM for fNIRS approach was developed based on effective connectivity among approximate sensor level regions near the cortical surface and validated in two illustrative participants - one adult (Tak et al., 2015) and one infant (Bulgarelli et al., 2018). To overcome this limitation, we propose combining diffuse optical tomography (DOT) with DCM, enabling us to derive precise source level priors for neuronal sources beneath the pial surface and incorporate them into the DCM for fNIRS pipeline. We validate the proposed method using a cohort of more than 100 participants.

DOT reconstructs voxelwise hemoglobin signals at the source level beneath the cortex from sensor level optical density measurements by inverting a forward light propagation model (Bluestone et al., 2001; Boas et al., 2004; Custo et al., 2010; Joseph et al., 2006; Wheelock et al., 2019). Compared with conventional fNIRS topography, DOT offers improved depth sensitivity and spatial resolution, allowing brain derived activation to be distinguished from confounding signals originating in superficial tissues such as scalp vasculature or cerebrospinal fluid (CSF) (Boas et al., 2004; Custo et al., 2010; Hoshi, 2016). This improvement is achieved by incorporating anatomical priors from either subject specific structural MRI or a standardized head template when individual MRI data are unavailable (Custo et al., 2010, Hoshi, 2016; Wheelock et al., 2019).

Previous studies have inferred causal networks from DOT data using psychophysiological interaction (PPI) analysis (Hassanpour et al., 2017, Piva et al., 2017; Ávila-Sansores et al., 2021). Unlike DCM, however, PPI operates directly on measured hemodynamic time series, limiting causal inference to the temporal resolution of those measurements (Stephan and Friston, 2010). Here, we propose a method for estimating effective connectivity with DCM using source locations identified from DOT reconstructed signals. We apply this pipeline to a Go/No-Go experimental dataset, a standard paradigm for probing inhibitor control. We hypothesize that the improved depth sensitivity and spatial precision of DOT will yield more accurate neuronal source localization, thereby providing superior effective connectivity estimates relative to traditional sensor-level DCM analysis of fNIRS data.

## 2. Methods

The analysis pipeline consists of three components: (i) reconstruction of DOT images by solving the inverse problem of the forward light model, (ii) preprocessing and statistical parametric mapping (SPM) of the reconstructed voxel-wise DOT signals to identify brain regions activated by the experimental protocol, enabling source-level inference of brain activation, and (iii) application of DCM to optical density measurements at the identified source locations. A schematic overview of the proposed method is provided in Fig. 1.

**Figure 1.**
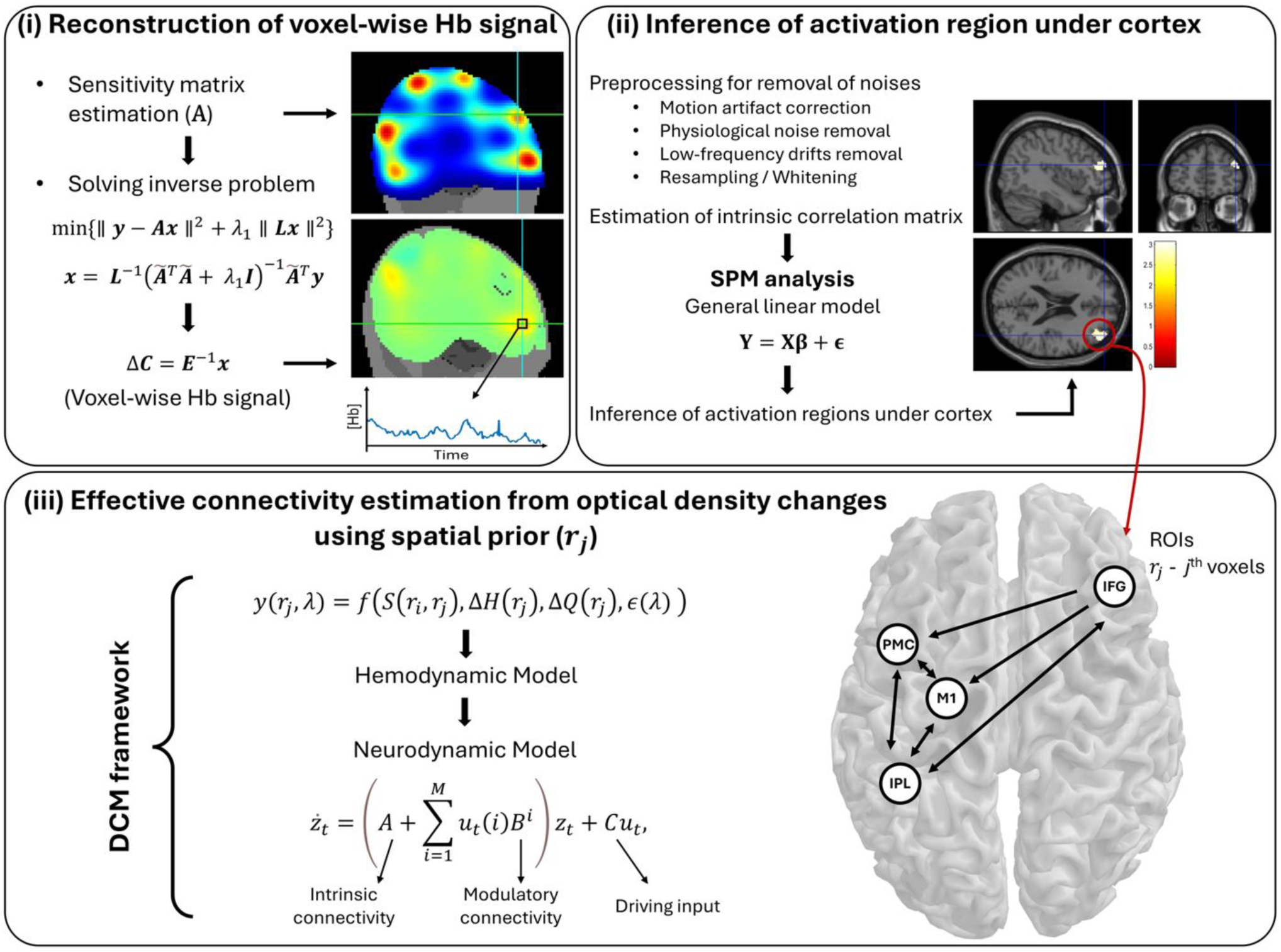
Schematic of the analysis pipelines. The procedures comprises three steps: (i) reconstruction of diffuse optical tomography (DOT) images, by solving the inverse problem of the forward light model, resulting in voxel-wise hemoglobin signals, (ii) source level inference of brain activation via preprocessing and statistical parametric mapping (SPM) of the reconstructed voxel-wise DOT signals, and (iii) estimation of effective connectivity between the identified source locations from optical density changes using dynamic casual modelling (DCM).

### 2.1. Reconstruction of Diffuse Optical Tomography

DOT reconstruction was performed in two major steps (Eggebrecht and Culver, 2019; Speh et al., 2025): (i) generation of the forward light model and (ii) image reconstruction by solving the inverse problem. The forward light model describes the relationship between surface level optical measurements and changes in optical properties - specifically, concentrations of HbO and HbR - within the volume. This relationship can be expressed as follows:

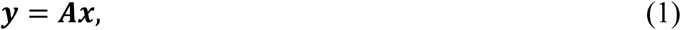

with ***y*** represents the optical measurements from the fNIRS channels, ***x*** denotes the voxel-wise variations in absorption and scattering corresponding to hemodynamic changes, and ***A*** is the sensitivity matrix that characterizes the spatial sensitivity of each channel to absorption and scattering changes. The sensitivity matrix is constructed from the forward light model, which is derived from the Radiative Transport Equation (RTE) (Wheelock et al., 2019). An approximate solution to the RTE was computed using the NIRFAST package (Dehghani et al., 2009), which employs a fast finite element modeling (FEM) routine. Computing the sensitivity matrix requires information about the tissue boundary geometry and spatial distribution of optical properties (the head model). We used tissue segmentation of the Montreal Neurological Institute (MNI) 152 standard atlas (Eggebrecht and Culver, 2019) when estimating the sensitivity matrix (Wheelock et al., 2019; Custo et al., 2010, Ferradal et al., 2014).

With the forward light model specified, DOT reconstruction was performed by solving the following minimization problem using the NeuroDOT toolbox (Eggebrecht and Culver, 2019; Speh et al., 2025):

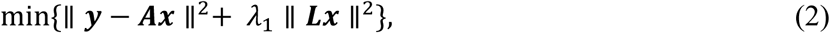

where the term λ_1_ ∥ ***Lx*** ∥^2^ serves as a penalty for image variance. This includes a spatially variant regularization term λ_2_, incorporated into the diagonal of *L* as follows:

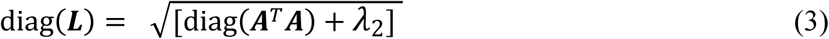

The solution of Eq. (3) was then obtained using the Moore-Penrose generalized inverse:

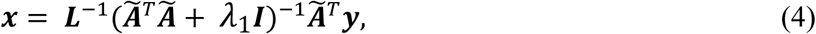

where **Ã**= ***AL***^−**1**^, and ***I*** is the identity matrix.

This gave the relative changes in oxy- and deoxy-hemoglobin concentrations (Δ***C***) as follows

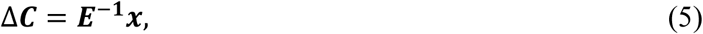

where ***E*** is the extinction coefficient matrix, and Δ***C*** is the matrix representing concentration changes in HbO and HbR.

Regularization parameters, λ_1_and λ_2_ in Eqs. (2) and (3) significantly influence the characteristics of the reconstructed image. The Tikhonov regularization parameter λ_1_controls the degree of spatial smoothing: a low value enhances high-frequency spatial details (including noise), while a high value yields a smoother image (Culver et al., 2001; Wheelock et al., 2019). The spatially variant regularization parameter λ_2_ further improves image quality by reducing localization errors and enhancing resolution uniformity and contrast (Culver et al., 2003; Wheelock et al., 2019). Importantly, λ_2_ also enables depth-dependent tuning of the reconstruction, with smaller values allowing deeper imaging below the surface and larger values constraining reconstruction to shallower regions (Wheelock et al., 2019). The optimal settings of λ_1_ and λ_2_ depend on the geometry and characteristics of the imaging system. In our study, we used λ_1_ = 0.01 and λ_2_ = 0.1.

### 2.2. Statistical Parametric Mapping for Diffuse Optical Tomography

After reconstructing the DOT images with Eqs. (1) - (5), we applied a general linear model (GLM) at both within-subject and group levels to identify brain regions activated by the experimental protocol (Friston et al., 1994; Hassanpour et al., 2014; Ye et al., 2009; Tak et al., 2016)

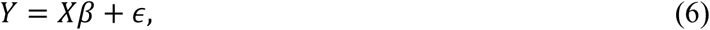

where *Y* is an [*M* × *N*] matrix of HbO or HbR concentration changes (Δ***C***); M is the total number of time points (scans) and N is the total number of voxels; *X* is an [*M* × *L*] desgin matrix containing *L* regressors of interest and nuisance covariates; *β* is an [*L* × *N*] matrix of regression coefficients; and *ϵ* is an [*M* × *N*] matrix of zero-mean, normally distributed errors with covariance *σ*^2^***V***, where *σ*^2^ is voxel-specific variance and ***V*** is temporal autocorrelation matrix common to all voxels.

For voxel-specific hemoglobin responses (*Y*), we applied a preprocessing pipeline to remove confounding effects such as motion artifacts and physiological noise from the time series. Specifically, the reconstructed HbO signals were preprocessed using the following steps: (i) motion artifact correction based on moving standard deviation and spline interpolation (Scholkmann et al., 2010), (ii) removal of physiological noise (respiration and cardiac pulsation) using an infinite impulse response (IIR) Butterworth filter with stopband frequencies of 0.12−0.35 Hz and 0.7−2.0 Hz, and (iii) removal of very low-frequency drifts using a high-pass filter based on a discrete cosine transform (DCT) set with a cutoff frequency of 1/64 Hz. The preprocessed data were then resampled to 1 Hz and temporally whitened using an autoregressive (AR) model of order 1.

We fitted a GLM model (see Eq. (6)) to the preprocessed hemoglobin signals to estimate subject-level effects. The design matrix comprised the task stimulus timings convolved with the canonical hemodynamic response function and its temporal and dispersion derivatives. Source space parameters (*β*) were estimated using generalized least square methods (Worsely and Friston 1995). Group-level activation maps were then obtained with a random-effects analysis based on summary statistics across subjects. Conventional statistical parametric mapping for fNIRS provides sensor-level inference-that is, activated region on the scalp surface. In contrast, our DOT-based SPM framework enables source-level inference, localizing activations at the voxel level beneath the cortex. The cortical activation patterns rendered by SPM-fNIRS and SPM-DOT were spatially consistent; however, the DOT approach generally produced higher statistical significance.

The group-level result was used to identify the regions of interest (ROIs) for the subsequent DCM analysis.

### 2.3. Dynamic causal modeling for fNIRS with DOT-based spatial priors

We extended the generative model of dynamic causal modeling for fNIRS (Tak et al., 2015) by more precisely specifying neuronal source locations *r*_*j*,ℎ_ statistically inferred from the DOT reconstruction (i.e., source-space analysis). The optical forward model in the DCM framework is expressed as

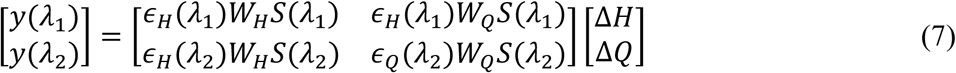

where *y*(λ_1_) and *y*(λ_2_) are [*N* × 1] vectors of optical-density changes measured at wavelengths λ_1_ and λ_2_, respectively; N denotes the number of channels; and Δ𝐻 and Δ𝑄 are [𝐽 × 1] vectors of voxel-wise HbO and HbR concentration changes, respectively. The *J* voxels of interest were defined from statistical inference of the DOT-reconstructed hemoglobin signals; 𝑆(λ) is the sensitivity matrix whose (*i*, *j*)*-* th element is given by 𝑆_𝑖,*j*_(λ) = 𝐺(*r*_𝑖,𝑠_, *r*_*j*,ℎ_)𝐺(*r*_*j*,ℎ_, *r*_𝑖,𝑑_)/ 𝐺(*r*_𝑖,𝑠_, *r*_𝑖,𝑑_) where *r*_𝑖,𝑠_ and *r*_𝑖,𝑑_ denote the source and detector positions for the i-th channel, and *r*_*j*,ℎ_is the hemodynamic source position of the *j*-th voxel; *ϵ*_𝐻_and *ϵ*_𝑄_ are the extinction coefficients of HbO and HbR; and 𝑊_𝐻_ and 𝑊_𝑄_are [*N* × *N*] correction matrices that compensate for pial-vein contamination.

This optics model was combined with the conventional generative models of hemodynamics and neuronal activity. The model parameters – including effective connectivity at the neuronal level – were estimated using a variational Bayesian inversion algorithm.

In order to statistically validate the performance of the proposed framework of optics model, we generated two models: one with hemodynamic source locations specified with SPM for DOT and one with hemodynamic source locations specified with SPM for fNIRS. The winning model was selected using Bayesian inference at the model level based on computed model evidence.

## 3. Experiments

### 3.1. Datasets

In this work, we used a dataset comprising 104 typically developing children (mean age = 8.05 years, standard deviation = 1.78; range: 6–14 years). The data were collected in previous studies (Sutoko et al., 2018, 2019; Tokuda et al., 2018), and here we apply the proposed method to estimate source-level regional activity and effective connectivity and to evaluate its performance. Detailed descriptions of the dataset have been presented in relevant literature (Sutoko et al., 2018, 2019; Tokuda et al., 2018). Below we summarize the experimental setup and protocol.

Written informed consent for participation was obtained from the parents of all subjects. The study was approved by the Ethics Committees of Jichi Medical University Hospital, and the International University of Health and Welfare, and conducted in accordance with the Declaration of Helsinki.

The multichannel fNIRS system ETG-4000 (Hitachi Medical Corporation, Kashiwa, Japan) was used to measure the subject’s hemodynamic activity using two wavelengths of near-infrared light (695 and 830nm). The fNIRS probes were positioned to cover the lateral prefrontal cortices and parietal lobes of both hemispheres. Specifically, for each hemisphere, a set of eight illuminating sources and seven detecting probes was arranged in a 3x5 probe holder configuration with an inter-probe distance of 3 cm, resulting in a total of 44 channels. The detailed system setup has been previously described in Monden et al., (2012, 2015). The positions of the sources, detectors probes, and channels were measured and transformed into corresponding positions in Montreal Neurological Institute (MNI) space using the NFRI functions (Okamoto et al., 2004; Singh et al., 2005), as shown in Fig. 2.

**Figure 2.**
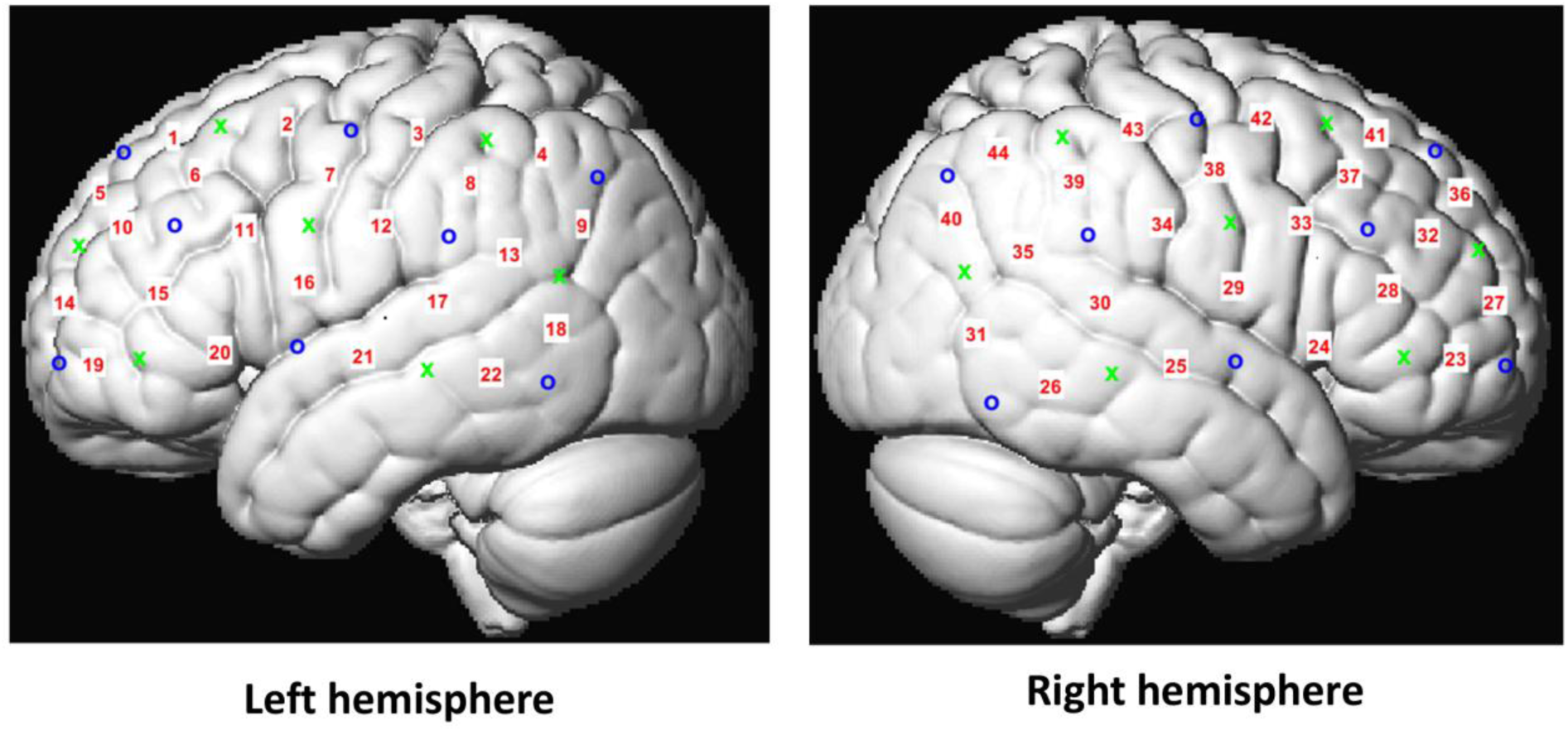
fNIRS acquisition setting. The fNIRS probes were placed to cover the lateral prefrontal cortices and parietal lobe of both hemispheres. For each hemisphere, a set of 8 illuminating sources and 7 detecting probes were arranged in a 3x5 probe holders with an inter-probe distance of 3 cm, which results in 44 channels. fNIRS: functional near-infrared spectroscopy.

The subjects were examined for hemodynamic changes related to response inhibition while performing a Go/Nogo task. As shown in Fig. 3, the task paradigm consisted of six blocks alternating between 24-second Go (baseline) and 24-second Nogo (target) trials. Before each trial, instructions were displayed for 3 seconds to inform the subjects about the upcoming condition. The total session lasted approximately 6 minutes. In the Go trials, subjects were presented with an alternating images of a lion and a giraffe and were instructed to press a button in response to both images. In the Nogo trials, a no-go image (tiger) was presented for 50% of the time, requiring subjects to either respond (Go stimulus) or inhibit their response (Nogo stimulus) for each half of the trial. The stimuli were presented at a frequency of 1Hz. All subjects practiced the task in advance to become familiar with the paradigms prior to measurement.

**Figure 3.**
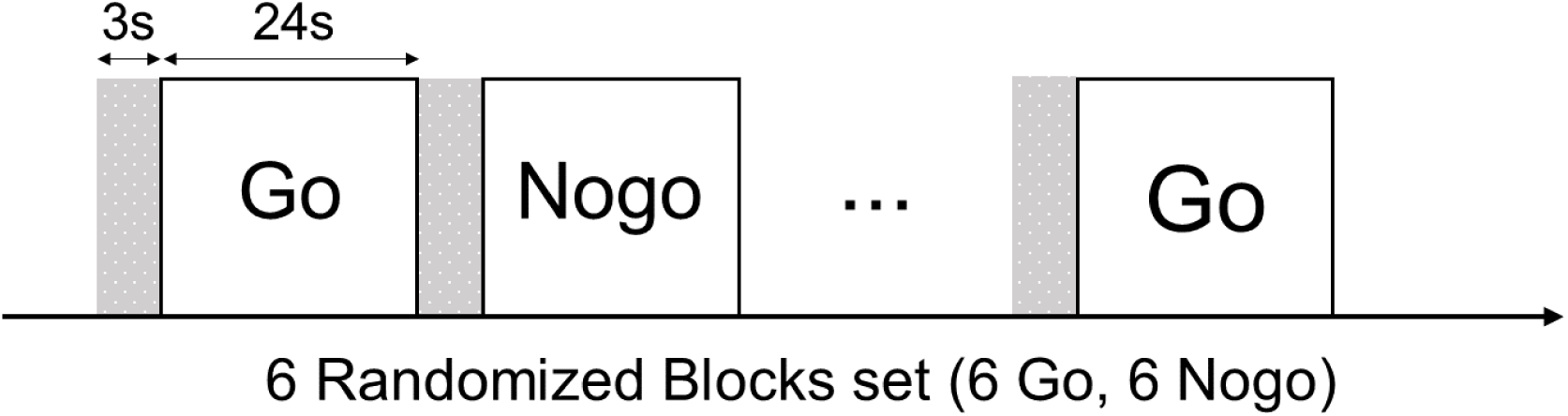
Schematic of the experimental paradigm. The task paradigm includes 6 block set that alternated between 24s Go (baseline) and 24s Nogo (target) trials. There are instructions displayed for 3s before each trial. The initial pre-scan resting state of 10s was also included.

Response inhibition is a core component of cognitive control that enables individuals to suppress inappropriate or unwanted actions (Simmonds et al., 2008; Aron, 2011). Deficits in this ability contribute to several neuropsychiatric disorders, most notably attention-deficit/hyperactivity disorder (ADHD), in which patients struggle to curb impulsive behaviour (Aron & Poldrack, 2005). The Go/No-Go task is the standard paradigm for probing response inhibition: participants press a button as quickly as possible when a Go cue appears and withhold the response when a No-Go cue appears. Although any Go-to-No-Go ratio can be used, many studies present Go cues more frequently (typically 70–80 % of trials) to create a strong pre-potent tendency to respond; successful withholding on the rarer No-Go trials thus becomes a sensitive measure of inhibitory control (Garavan et al., 1999; Rubia et al., 2001; Simmonds et al., 2008).

Neuroimaging work has consistently identified a right-lateralised activation pattern during No-Go trials, with the right inferior frontal gyrus (rIFG) emerging as a critical node (Garavan et al., 1999; Aron et al., 2004). However, these studies report undirected activation maps or functional (correlational) connectivity; they do not reveal directed (causal) influences between regions. Consequently, the precise pathways by which the rIFG suppresses motor output remain unclear.

To address this gap, we applied our DOT-informed DCM framework to estimate effective connectivity, directed causal influences, between the IFG and the motor network during the Go/No-Go task. By modelling how No-Go trials explicitly modulate these directed connections, our approach could provide the evidence of how the rIFG exerts top-down inhibitory control over motor regions.

### 3.2. Analysis: Connectivity Model for Experimental Data and Parameter Estimation

In this study, we evaluated three connectivity models (Fig. 4) to investigate how the right and/or left IFG influence the motor network comprising left primary motor cortex (M1), premotor cortex (PMC), and inferior parietal lobule (IPL) in suppressing motor responses during response inhibition. Effective connectivity among regions was modeled with the following system of parameter matrices:

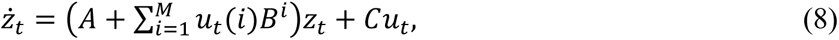

**Figure 4.**
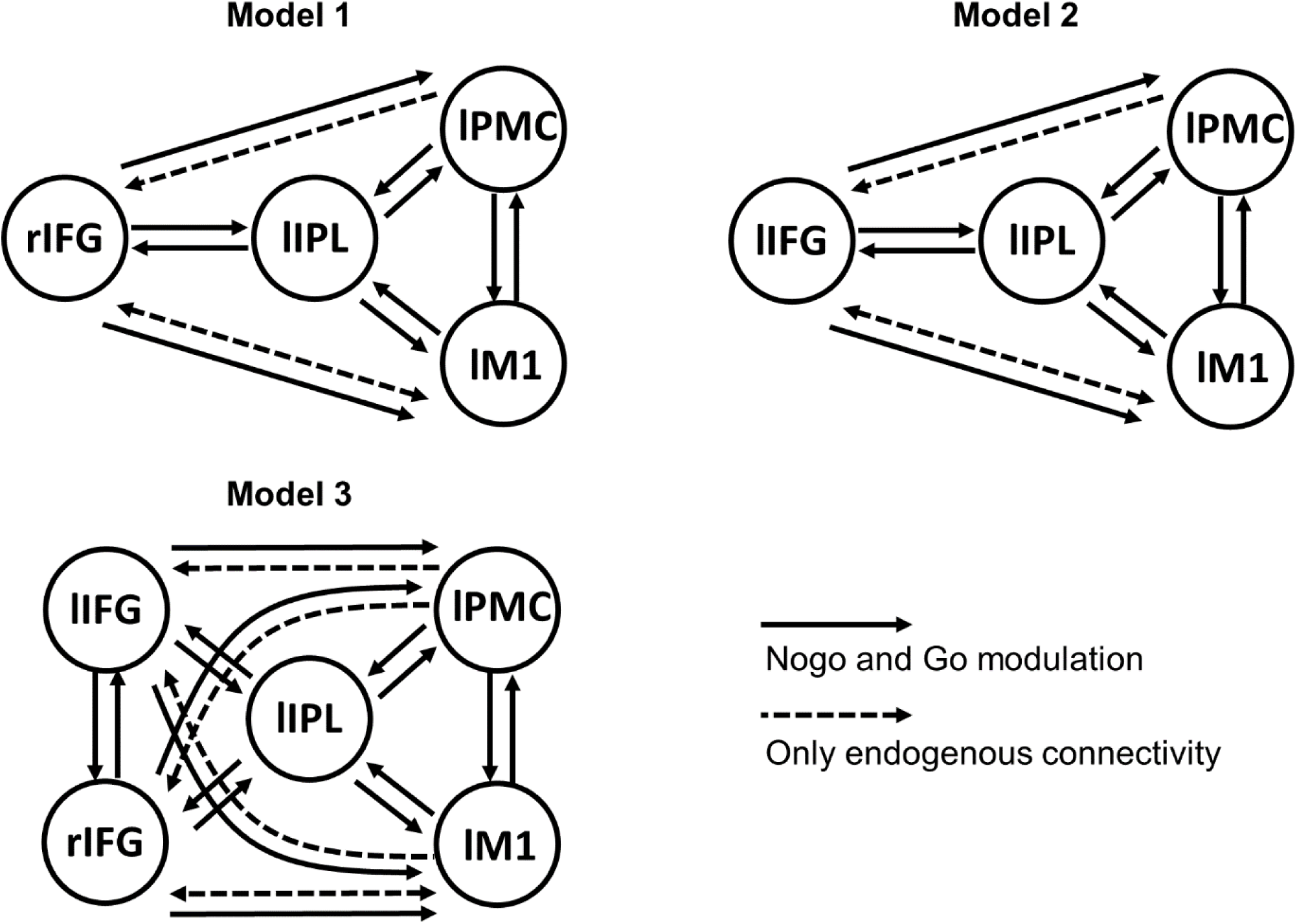
Models for dynamic causal modeling analysis for Go/Nogo task response inhibition. All models include the motor regions including the left primary motor cortex (lM1), left premotor cortex (lPMC), and left inferior parietal lobule (lIPL). The models are different in the exist of inferior frontal gyrus (IFG). Specifically, model 1 includes the right IFG (rIFG), model 2 includes the left IFG (lIFG), while model 3 includes both right and left IFG.

where 𝑧_𝑡_is the vector of neuronal activities at time *t*; its entry represents the activity of a distinct brain region. The matrix A encodes intrinsic, context-independent connectivity present in the absence of experimental manipulation, whereas each 𝐵^𝑖^captures task-dependent changes in connectivity associated with experimental condition i. The binary indicator 𝑢_𝑡_(𝑖) ∈ {0,1} specifies whether condition *i* is active at time *t*, and M denotes the total number of conditions. The matrix C defines the direct influence of experimental inputs on each region. The neuronal dynamics generated by this equation are subsequently passed to the hemodynamic model and then to the optical forward model, together forming the full generative framework of DCM for DOT.

In every model these motor regions are fully interconnected via endogenous (A-matrix) and modulatory (B-matrix) links, with the B-matrix capturing trial-specific modulation in both Go and Nogo conditions. Modulatory effects for the Go condition are encoded at the first index of the third dimension of the B matrix, whereas those for the Nogo condition are encoded at second index, allowing the corresponding parameters to be estimated separately.

The three models differ in their inclusion of the IFG regions. Model 1 incorporates the right IFG, which is reciprocally connected to the motor network (A matrix) and exerts modulatory influences on that network (B matrix). Because connections between the IPL and IFG are critical for the executive control of goal-directed and stimulus-driven attention-functions required for response selection-we specified reciprocal modulatory connectivity between these regions. Model 2 replaces the right IFG with the left IFG, while Model 3 contains both the right IFG and left IFG. In all three models, every region receives the task input through the C matrix.

After specifying the DCM connectivity models, we estimated their parameters - posterior distributions and log model evidence values - for each participant using the variational Laplace method. Bayesian model selection (BMS) with a random-effects approach was then applied to identify which of the three models best explained the observed data. To characterize group-level effective connectivity, we performed Bayesian model averaging (BMA) with Bayesian parameter averaging; the resulting posterior means represent group-level connection strengths. Parameter significance was assessed with one sample t-tests (*p* < 0.05). We additionally conducted paired t-tests comparing modulatory connectivity in Go versus Nogo trials to determine task-dependent changes among regions during response inhibition. This DCM pipeline allowed us to address two experimental questions: (i) which region-left IFG, right IFG, or both-drives response inhibition, and (ii) how these IFG regions influence the motor network to suppress motor responses.

## 4. Results

Fig. 5 shows the group-level activation maps derived from reconstructed DOT data for the Nogo > Go and Go > Nogo contrasts. Peak coordinates and statistical details are summarized in Table 1, using an uncorrected threshold of *p* < 0.005 (Lieberman et al., 2009). In the Nogo > Go contrast, significant activation was observed in the right IFG and right middle frontal cortex. In contrast, the Go > Nogo condition revealed an activation cluster spanning the left M1, PMC, and IPL, along with the left IFG.

**Figure 5.**
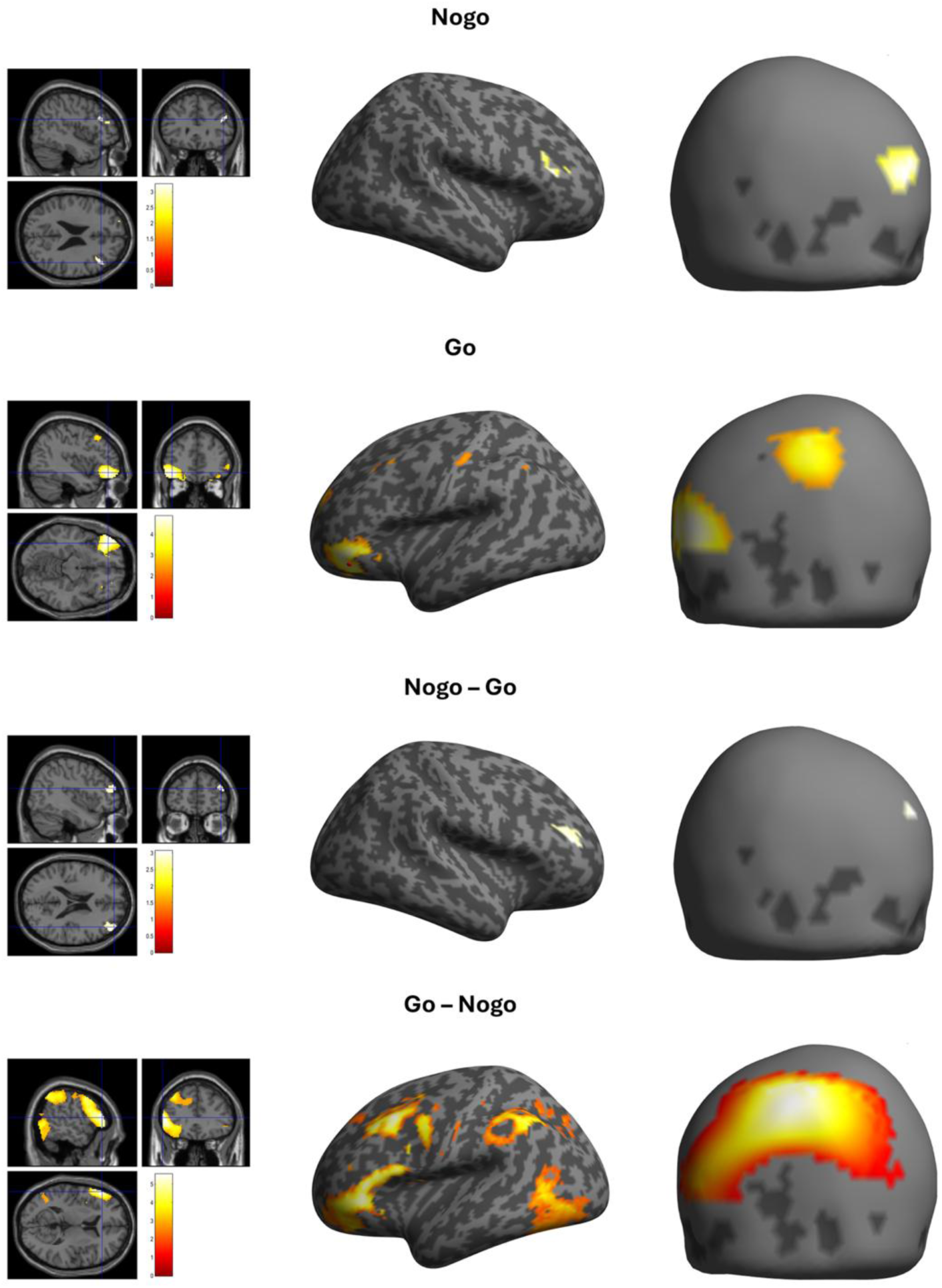
SPM for DOT reconstructed images results. Sectional view (1^st^ column) and render brain view (2^nd^ column) of the significant activated regions in the Nogo, Go, Nogo > Go, and Go > Nogo contrasts were shown, respectively. The SPM results were thresholded at uncorrected p < 0.005. The activation pattern was highly consistent in comparison with results of SPM for fNIRS (3^rd^ column). Details of the activated regions and statistics were summarized in Table 1. SPM: statistical parametric mapping, DOT: diffuse optical tomography, fNIRS: functional near-infrared spectroscopy

**Table 1.**
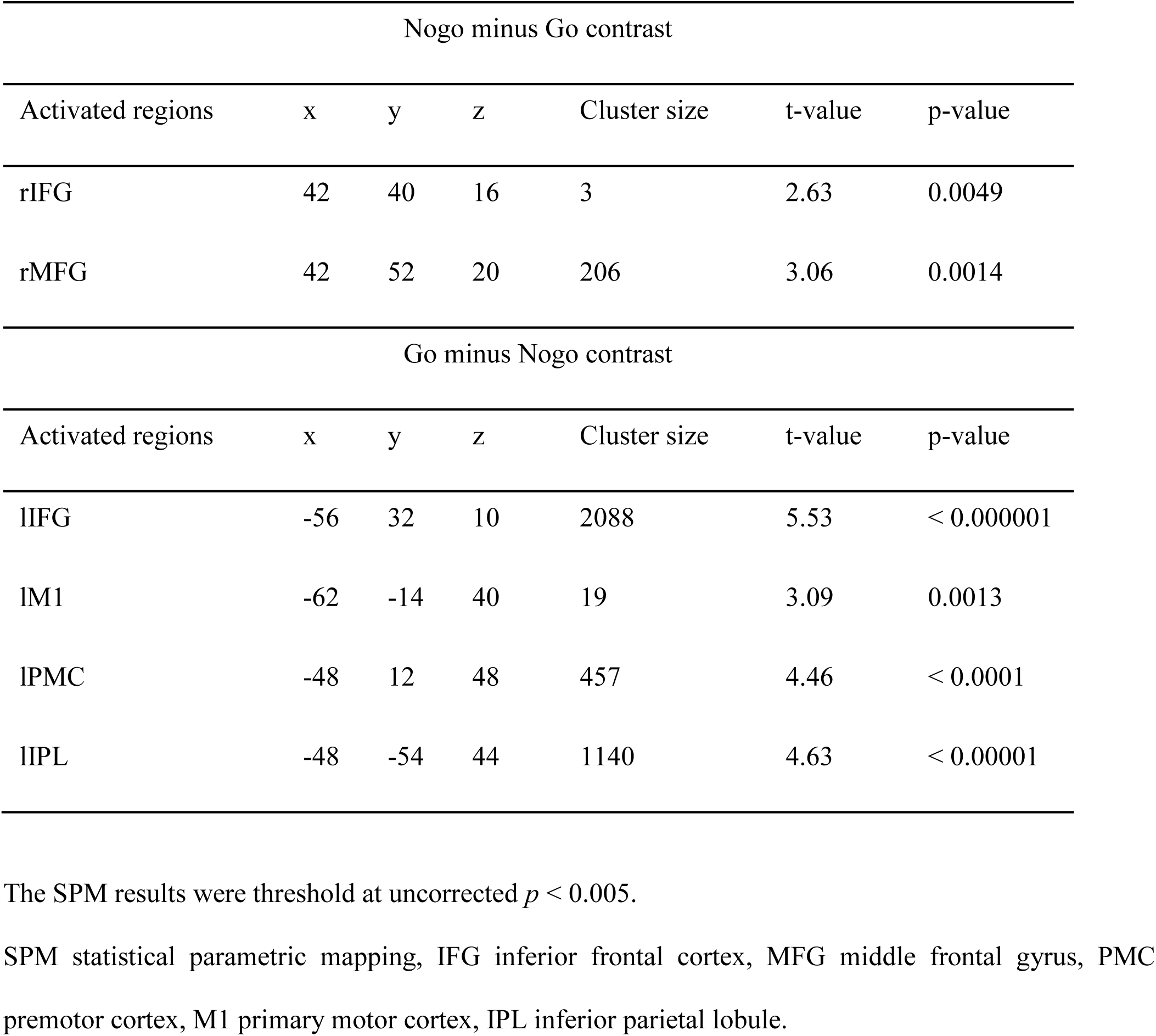
Group MNI coordinate of Region of interests for all subjects during Go/Nogo task.

| Nogo minus Go contrast |  |  |  |  |  |  |
| --- | --- | --- | --- | --- | --- | --- |
| Activated regions | x | y | z | Cluster size | t-value | p-value |
| rIFG | 42 | 40 | 16 | 3 | 2.63 | 0.0049 |
| rMFG | 42 | 52 | 20 | 206 | 3.06 | 0.0014 |
| Go minus Nogo contrast |  |  |  |  |  |  |
| Activated regions | x | y | z | Cluster size | t-value | p-value |
| lIFG | -56 | 32 | 10 | 2088 | 5.53 | < 0.000001 |
| lM1 | -62 | -14 | 40 | 19 | 3.09 | 0.0013 |
| lPMC | -48 | 12 | 48 | 457 | 4.46 | < 0.0001 |
| lIPL | -48 | -54 | 44 | 1140 | 4.63 | < 0.00001 |

The SPM results were threshold at uncorrected $p < 0.005$ .
SPM statistical parametric mapping, IFG inferior frontal cortex, MFG middle frontal gyrus, PMC premotor cortex, M1 primary motor cortex, IPL inferior parietal lobule.

These patterns were expected because actual hand movements occured exclusively in response to Go trials, resulting in greater activation of motor-related regions. Conversely, inhibitory control to suppress automatic motor responses, as well as executive attentional processes necessary for withholding responses, are functions commonly attributed to IFG activity and were more strongly engaged during Nogo trials, thus explaining the increased IFG activation. Indeed, previous fMRI and fNIRS studies (Garavan et al., 1999; Rubia et al., 2001; Simmonds et al., 2008) employing the Go/Nogo paradigm have consistently reported similar activation regions, undergoing the validity of our results. In addition, sensor-level SPM analysis of fNIRS data (Fig. 5 - 3^rd^ column) revealed highly consistent activation patterns with the DOT results, further confirming the reliability of the observed activations. The different functional roles of the right and left IFG regions in response inhibition are addressed in more detail in the Discussion section. These identified regions served as principal nodes within the motor network for subsequent DCM analysis.

Dynamic causal modeling (DCM) was fitted to the optical density signals, and directed connectivity among specified network nodes, whose cortical locations were identified using the SPM for DOT approach, was subsequently estimated. Fig. 6 presents the Bayesian model selection results comparing the statistical significance of DCM parameters derived from DOT node locations (source level analysis) with those from conventional fNIRS node locations (sensor level analysis). As illustrated in the first column of Fig. 5, SPM for DOT identifies activated regions located deeper beneath the cortical surface. In contrast, due to its reliance on sensor level analysis, SPM for fNIRS primarily identifies activations on or near the cortical surface, as shown in the third column of Fig. 5. Leveraging this property,

**Figure 6.**
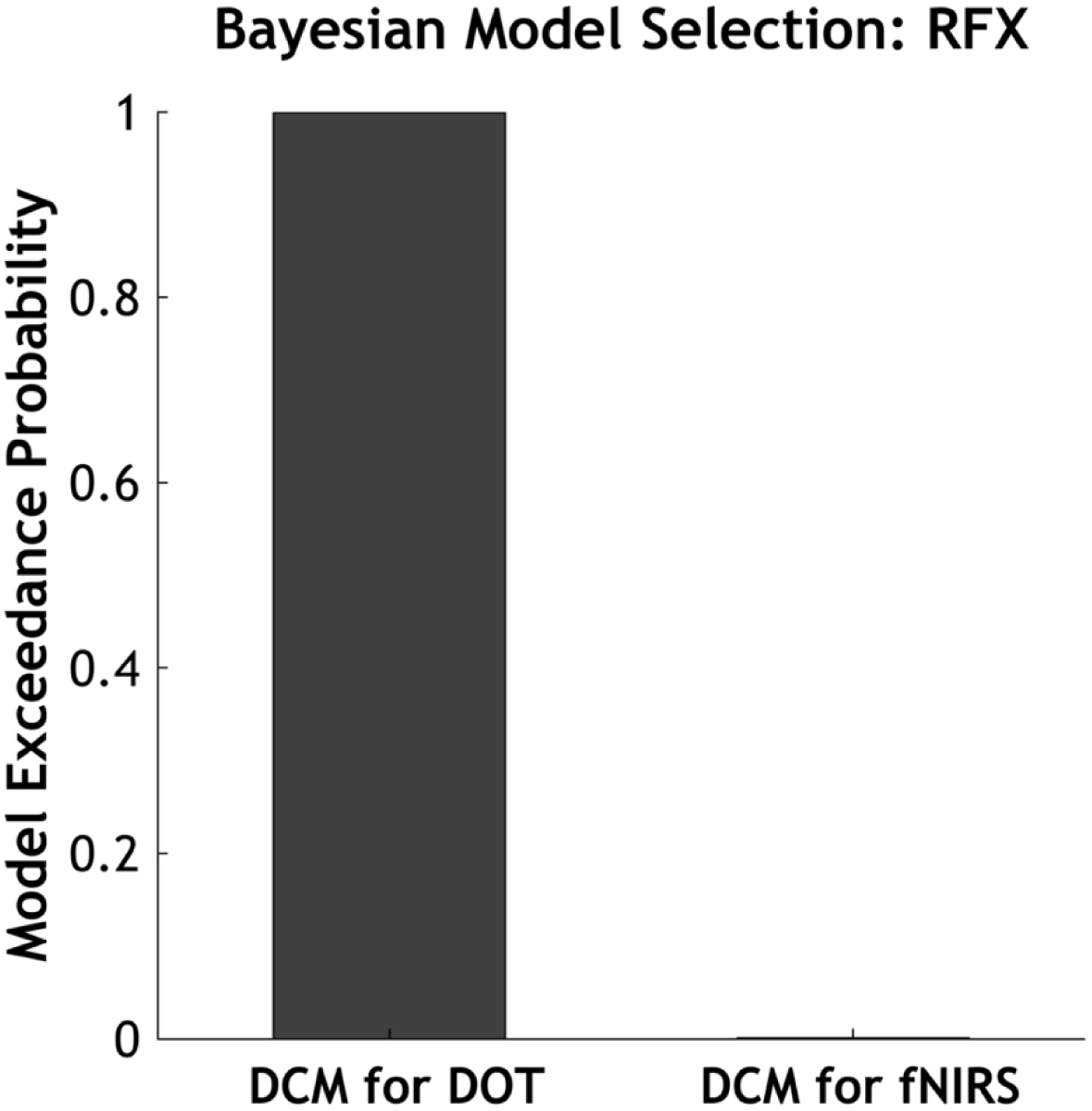
Bayesian model selection based on random-effects analysis (RFX) results for comparing model estimated using DOT node locations (source level analysis) or conventional fNIRS node locations (sensor level analysis). The results show that the proposed source level approach (DCM for DOT) (Model 1) outperformed the sensor level approach (DCM for fNIRS). DOT diffuse optical tomography, DCM dynamic causal modelling, fNIRS functional near-infrared spectroscopy.

Bayesian inference at the model level (Fig. 6) clearly indicated that the proposed source level approach (DCM for DOT) outperformed the sensor level approach (DCM for fNIRS).

Consequently, in subsequent analyses, we exclusively utilized the source level DCM for DOT method to identify the optimal network model explaining our experimental data. Bayesian model selection using random effects analysis revealed that model 1 was the most appropriate, exhibiting the highest exceedance probability (Fig. 7). This model specifically incorporates the right IFG region and its interactions with the motor network that comprises of PMC, IPL and M1 regions.

**Figure 7.**
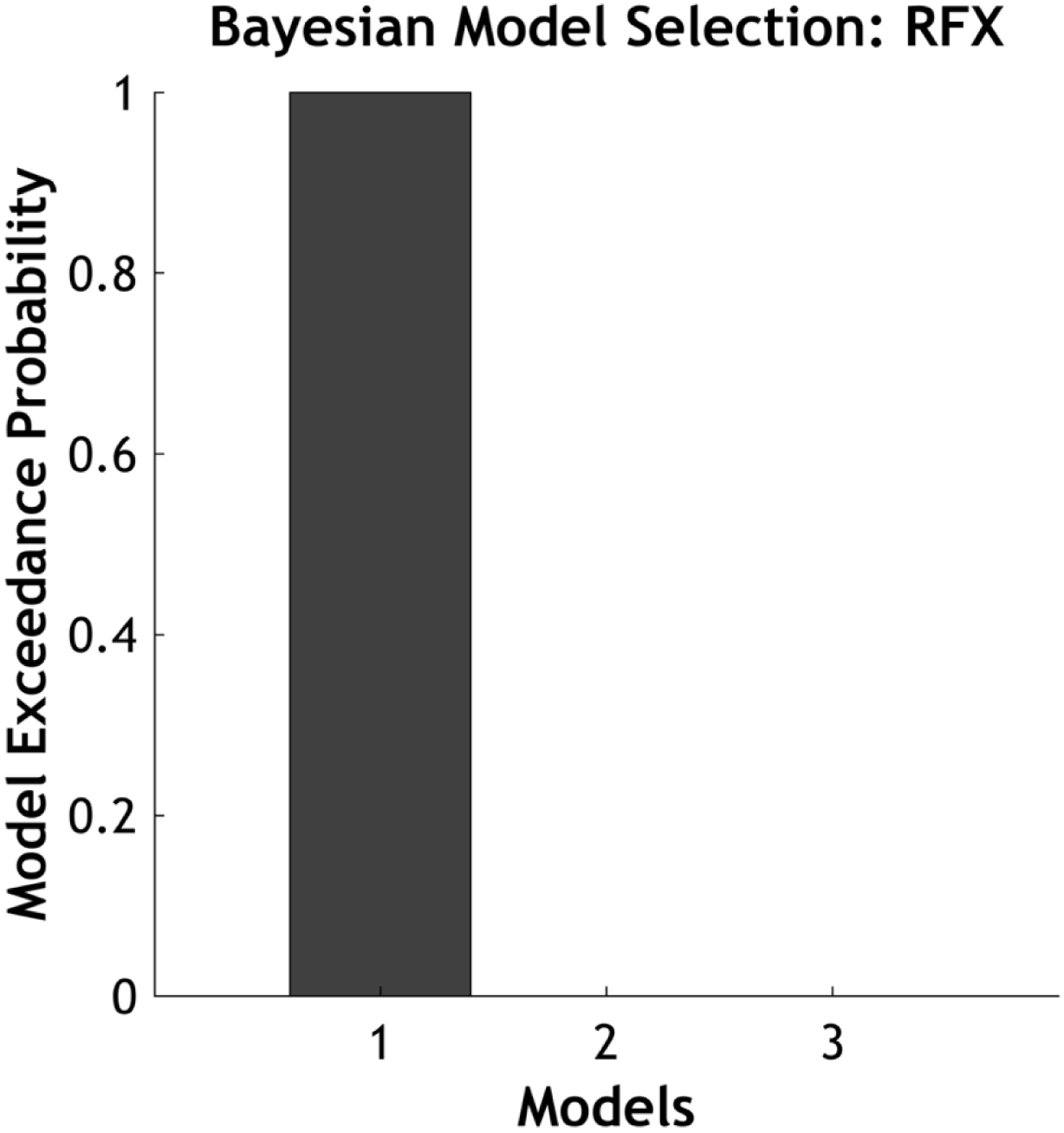
Bayesian model selection based on random-effects analysis (RFX) result. The winning model structure that best explained the data is model 1 (which includes the right IFG region and its interactions with the motor network that comprises of PMC, IPL and M1 regions), showing the highest model exceedance probability. IFG inferior frontal gyrus, PMC premotor cortex, IPL inferior parietal lobule, M1 primary motor cortex.

The posterior means of endogenous and modulatory connectivity parameters for the winning model (Model 1) were estimated using Bayesian model averaging, and their corresponding values are presented in Figure 8. Significant posterior parameter estimates that survived a statistical threshold of *p* < 0.05 are indicated by an asterisk (*). Positive connectivity parameters imply that an increase in activity in one region leads to an increase rate of activity change in another region, and vice versa. Connectivity from the right IFG (rIFG) to the left M1 (lM1) was significantly negative during Nogo modulation (-0.05). Furthermore, the Nogo-minus-Go contrast revealed significantly stronger inhibitory effects from the rIFG to lM1 during Nogo compared to Go trials (effective connectivity difference: -0.07). Within the motor network, overall connectivity between regions decreased during Nogo relative to Go trials. Specifically, significant connectivity reductions were observed between left IPL (lIPL) and left M1 (lM1) (lIPL → lM1: -0.07; lM1 → lIPL: -0.07), and between lM1 and lPMC (PMC → lM1: -0.06; lM1 → lPMC: -0.07). It is noted that the decrease in connectivity from the rIFG to the left M1 in the Nogo-minus-Go contrast is not similar to the reduction in connectivity between motor regions and thus, is not due to the overall decrease in connectivity. The connectivity among motor regions showed significant positive connectivity in the Go contrast and less significant in the Nogo contrast, while the connectivity from the rIFG to left M1 demonstrated the opposite pattern. Additionally, not all connectivity from the rIFG was decreased in the Nogo-minus-Go contrast, as increased connectivity was observed from the right IFG to the left IPL, though this increase did not reach statistical significance.

**Figure 8.**
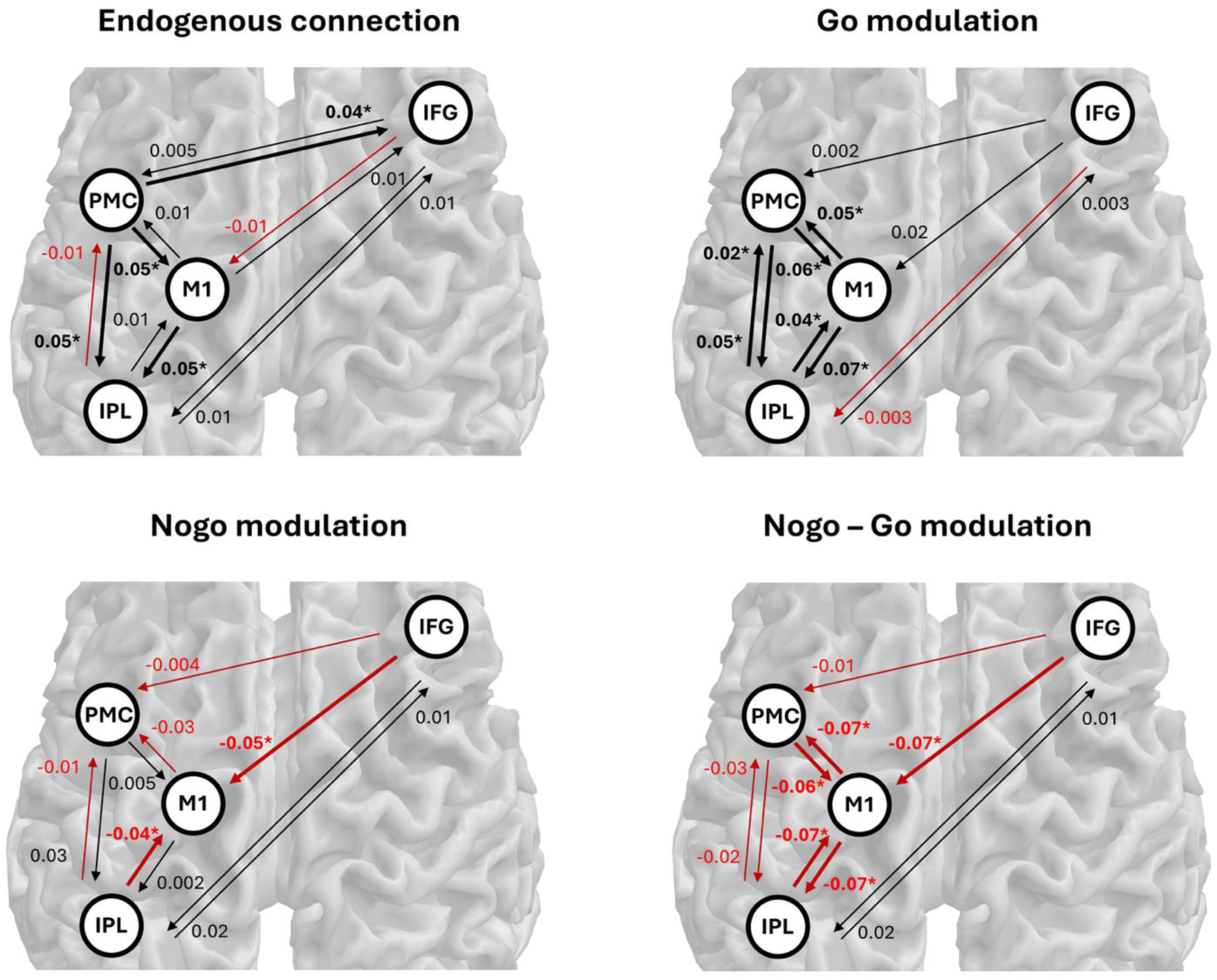
The group-level posterior mean of the endogenous and modulatory connectivity for the model 1, which is the winning model including the rIFG. There is a significant inhibitory connectivity from the rIFG to the lM1 during Nogo trials, and this connectivity is significantly decreased in the Nogo-minus-Go contrast. This suggested a role of rIFG in response inhibition control process. * indicates significant connectivity with t-test threshold *p* < 0.05. rIFG right inferior frontal gyrus, lPMC left premotor cortex, lIPL left inferior parietal lobule, lM1 left primary motor cortex.

## 5. Discussion

The main contribution of this paper is incorporating the identification of activated regions at the hemodynamic source level into causal network analysis for fNIRS signals. This was achieved by establishing a robust SPM analysis framework for reconstructed DOT signals and integrating its inference into the DCM-fNIRS analysis pipeline. By applying Bayesian model selection to an extensive experimental dataset, we demonstrated that our proposed method—DCM-fNIRS with precise source-level localization of hemodynamic activation—is statistically more powerful and preferable compared to the traditional approach based on approximate sensor-level locations.

Another key contribution is the identification of inhibitory control exerted by the right IFG on motor behavior, interpreted as an attentional mechanism from a motor network perspective. This finding was supported by applying the proposed method to a relatively large Go/Nogo task dataset, ensuring higher statistical power and reliability (Button et al., 2013). These results highlight the potential of fNIRS as a portable neuroimaging modality that, unlike MRI, can be readily deployed in naturalistic settings. This approach may be particularly useful in assessing children with ADHD (Monden et al., 2013, 2015; Poliakova et al., 2023), facilitating detection of imbalanced attentional control, and potentially informing therapeutic interventions aimed at restoring these cognitive functions.

In the following, we interpret the analysis results, focusing on effective connectivity derived using the proposed method and experimental dataset, and address its validity from a neuroscientific perspective. During the Nogo task, activated regions were found in the right inferior frontal cortex and middle frontal cortex. The results from SPM analyses of reconstructed DOT data strongly overlapped with those obtained using conventional sensor-level SPM for fNIRS, while additionally providing depth information about activation locations. This activation pattern is consistent with previous neuroimaging studies of response inhibition, which has shown that a right-lateralized network, especially involving the right IFG, is associated with inhibitory control (Garavan et al., 1999; Aron et al., 2004, Simmonds et al., 2008). The right IFG has been proposed as crucial for response inhibition, as damage to this region is frequently associated with impaired performance on inhibitory tasks (Aron et al., 2003). Furthermore, structural and functional MRI evidence from ADHD patients, who typically exhibit impaired response inhibition, suggests a relationship between deficits in the right IFG and compromised inhibitory control (Aron and Poldrack, 2005). Similarly, fNIRS-based studies have used activation in the right IFG to classify ADHD children from typically developing peers (Monden et al., 2015), and as a biomarker to assess clinical drug efficacy in treating ADHD (Monden et al., 2012). Our results thus demonstrate that the proposed pipeline successfully reconstructs activation maps with depth information directly from the fNIRS data.

In this study, dynamic causal modelling combined with Bayesian model selection determined that inhibitory control from the right IFG to motor network provided the best explanation of the experimental data, compared to models involving either the left IFG or both left/right IFGs. Specifically, the right IFG exerted a negative influence on the left M1 region both in the endogenous state and significantly during Nogo task modulation. Aron et al. (2004) previously proposed that the right IFG might suppress irrelevant response either indirectly through subcortical regions such as the subthalamus nucleus and brainstem, or directly through interactions with the motor cortex. Recent fMRI studies employing DCM in Go/Nogo tasks provide additional evidence of causal interactions between the right IFG and subcortical regions, reinforcing its critical role in response inhibition (Zhuang et al., 2023). Specifically, Zhuang et al. (2023) observed increased causal influences from the right IFG to the right caudate and thalamus, highlighting its central regulatory function in subcortical inhibitory control. Our findings further extend this view, demonstrating that the right IFG can also directly influence motor responses by exerting inhibitory control over motor cortical regions, particularly the primary motor cortex.

Our results also revealed distinct roles for the right and left IFG during the Go/Nogo task, suggesting hemispheric asymmetry and emphasizing the importance of right IFG in response inhibition. Specifically, unlike the right IFG, the left IFG did not exert significant inhibitory connectivity toward the left M1 during Nogo trials, but instead showed significant positive connectivity with the motor network region during Go trials. This implies that the left IFG may not play a central role in response inhibition. Previous studies have proposed that the left IFG is more involved in processes such as maintaining task or goal sets, rather than directly suppressing motor responses (Aron et al., 2004; Goghari et al., 2009). Supporting this, a recent study found that applying transcranial direct current stimulation (tDCS) to the left IFG did not enhance inhibitory control in healthy participants (Schroeder et al., 2022). These findings further support the right-lateralization of response inhibition and suggest that the left IFG does not have a critical role in this function.

The fNIRS approach has proven feasible in real-world clinical settings (Wheelock et al., 2019), making it well suited to studies involving young children and infants (Ferradal et al., 1991; White et al., 2012), and especially to investigations of children with ADHD (Gu et al., 2017; Monden et al., 2015). Consequently, our method could be extended to these populations to examine alterations in effective connectivity, and to elucidate the neural mechanisms underlying these disorders.

## 6. Conclusion

We have extended a DOT-informed DCM pipeline that incorporates source-level priors inferred from DOT reconstructed images into fNIRS effective connectivity analysis. Based on the data of children performing Go/Nogo tasks, we have estimated the effective connectivity among the IFG regions and the motor network. We have shown that our proposed method is more powerful and preferable compared to the traditional approach based on sensor-level location. Moreover, the estimated connectivity results suggested a crucial role of the right IFG in controlling motor response inhibition via directly influencing the primary motor cortex. This is highly consistent with previous studies and proves the validity of our novel approach. Therefore, this approach can be a promising method to study effective connectivity in young children and clinical settings.

## Data and Code Availability Statement

The data and code that support the findings of this study are available upon reasonable request.

## Acknowledgements

This work was supported by a grant of National Research Foundation of Korea (NRF) grant funded by the Korea Government (MSIT) (2022R1F1A1074729), grants from the Korea Basic Science Institute (C523400, D537150, C512130), and the National Research Council of Science & Technology(NST) grant by the Korea government (MSIT) (GTL25071-000).

## Author contributions

Truc Chu: Conceptulization, Formal Analysis, Investigation, Visualization, Validation, Software, Writing Kiyomitsu Niioka: Formal Analysis, Data Curation, Investigation, Writing Ippetia Dan: Resources, Data Curation, Investigation, Supervision, Project Administration, Writing Sungho Tak: Conceptualization, Methodology, Investigation, Validation, Software, Resources, Supervision, Project Administration, Funding Acquisition, Writing

